# APEX: Automated Protein EXpression in *Escherichia coli*

**DOI:** 10.1101/2024.08.13.607171

**Authors:** Martyna Kasprzyk, Michael A. Herrera, Giovanni Stracquadanio

## Abstract

Heterologous protein expression is an indispensable strategy for generating recombinant proteins. *Escherichia coli* (*E. coli*) is the most widely used microbial host for recombinant protein production due to its rapid growth, well-characterised genetics, and ability to produce recombinant proteins in high yields using modern recombinant DNA technology. However, while there is a plethora of robust protein expression protocols for *E. coli*, these methods are often unsuitable for high-throughput screening due to their significant resource and time consumption; these protocols are also susceptible to operator error and inconsistency.

To address these challenges, we have developed Automated Protein EXpression (APEX), a robust and automated protocol for recombinant protein production in *E. coli*. APEX leverages the accessible, open-source Opentrons OT-2 platform to automate microbial handling and protein expression with high precision and reproducibility. APEX can be configured to perform heat shock transformation, colony selection, colony sampling, microculturing and protein expression induction using a low-cost, minimal OT-2 hardware setup. We further demonstrate the efficacy of our automated transformation workflows using a variety of plasmids (2.7-17.7 kb), and exemplify the automated heterologous expression of a diverse array of proteins (27-222 kDa). Designed with customisation, modularity and user-friendliness in mind, APEX can be easily adapted to the operator’s needs without requiring any coding expertise.

APEX is available at https://github.com/stracquadaniolab/apex-nf under the AGPL3 license.

## Introduction

Recombinant protein production is a widespread and important technique for the study, screening and characterisation of proteins. Microbial cell factories are commonly used as recombinant expression systems, particularly *Escherichia coli* (*E. coli*), due to rapid growth, straightforward culturing conditions, well-understood physiology, easy transformation, and low costs [1]. Recombinant proteins have significant applications, including the production of high-value chemicals [2] and therapeutic proteins [3]. However, when processing numerous protein targets, such as in protein engineering where rapid identification of desirable properties is essential, these experiments become labour-intensive and inefficient, creating significant bottlenecks in research workflows [4]. Automation can transform these processes into high-throughput operations, significantly accelerating the processing of numerous protein targets.

Many workflows in biological laboratories have been successfully automated, including DNA assembly [5], sample preparation for next-generation sequencing [6], construction and high-throughput screening of peptide libraries [7], cell culturing and protein purification [8]. The adoption of automation has played a significant role across various fields, including drug discovery [9], clinical diagnostics [10], genomics [11], and proteomics [12]. Automation enhances efficiency by streamlining repetitive tasks [13], while improving reproducibility by mitigating human error, such as inconsistent pipetting [14]. By standardising processes, automation allows for consistent performance, regardless of operator experience, increasing both experimental success rates and the reliability of results.

While automation offers numerous advantages, the high costs often limit its accessibility to small and medium-sized laboratories. The Opentrons liquid handling robot addresses this challenge by providing an open-source platform that enables affordable laboratory automation. This platform has been successfully applied across various research fields, including DNA assembly methods such as Golden Gate [15] and BASIC [16], RNA extraction and RT-PCR diagnostics for SARS-CoV-2 [17, 18], protein purification and characterisation for enzyme discovery [19], and proteomics workflows for isobaric tag sample preparation [20] and shotgun proteomics [21].

Here, we developed an end-to-end automation pipeline for recombinant protein expression in *E. coli* called Automated Protein EXpression (APEX). This modular pipeline includes protocols for heat shock transformations, colony selection via spotting, colony sampling, and protein expression. It uses spreadsheet-based protocol configuration, allowing for customisation without programming experience. Protocols are generated via a computational pipeline that outputs a Python script (which can be directly uploaded to the Opentrons App) and a PDF file containing detailed protocol instructions and labware layouts for the reagents. To enhance reproducibility and ensure consistent generation of APEX protocols, we implemented the pipeline using Nextflow [22]. All necessary dependencies are packaged in a Docker container [23], allowing the pipeline to run seamlessly across various systems while providing a consistent environment for protocol generation.

We validated our APEX platform and assessed its usability through an extensive series of experiments. Initially, we miniaturised the transformation volumes in DH5*α E. coli* to minimise reagent use. Subsequently, we transformed using plasmids ranging from 2.7 to 17.7 kb and compared the performance of APEX with that of a human operator. We evaluated the pipeline’s ability for colony sampling, microculturing, and protein expression induction in BL21(DE3) *E. coli* using super folder Green Fluorescent Protein (sfGFP) as a benchmark. Finally, we executed the full pipeline with targets spanning molecular weights from 29 to 222 kDa, performing parallel runs with APEX and a human operator to compare their protein production outcomes. Collectively, our results demonstrate that APEX achieves robust and reproducible performance, ensuring high throughput while minimising reagents use and maximising efficiency of protein production workflows.

## Results

### The APEX workflow and implementation

The APEX workflow provides experimental scientists with a validated, optimised protein production protocol that leverages affordable lab automation. Crucially, it requires minimal computational experience for its adoption (Fig. 1a). We developed the protocols for the Opentrons OT-2 liquid handling robot, the most widely adopted low-cost, open-source system capable of protein expression. To ensure broad accessibility, APEX operates on a minimal required OT-2 setup, requiring only the thermocycler module and any combination of single or multi-channel pipettes.

**Figure 1:**
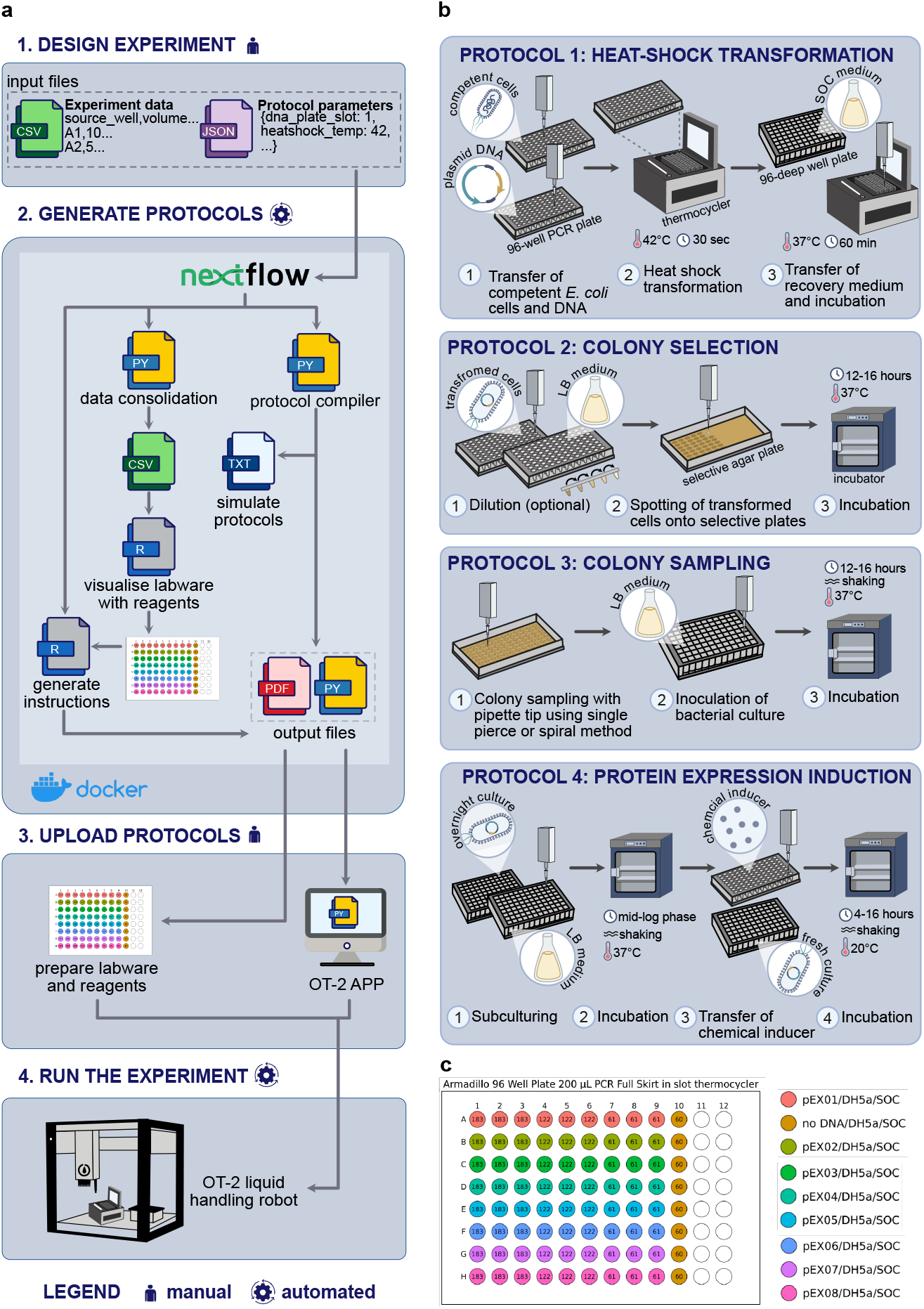
Automated Protein EXpression (APEX) workflow. **a**. The pipeline consists of four main steps: (1) User designs experiments using two input files: a JSON file for hardware and labware configuration, and a CSV file specifying experimental parameters such as source wells and volumes. (2) These input files serve three purposes: (i) generation of ready-to-use Python protocols from master templates, requiring no coding; (ii) generation of labware visualisation plots through R scripts; and (iii) generation of detailed user instructions in PDF format. The entire workflow is containerised in Docker and deployed as a Nextflow pipeline to ensure reproducibility and ease of use. (3) User uploads the Python protocols to the OT-2 app and prepares the robot deck. (4) Opentrons OT-2 robot executes the experiment, completing the workflow. **b**. APEX pipeline mirrors a typical protein expression workflow in *E. coli* through four modular protocols: Protocol 1: Heat shock transformation of DNA into cells; Protocol 2: Selection of transformed colonies on antibiotic-supplemented agar plates; Protocol 3: Colony sampling and cell culturing; Protocol 4: Protein expression induction. **c**. Example of visualised labware included in the instructions for user guidance geenrated by APEX.

The APEX pipeline automates the standard four-step protein expression process in *E. coli*: (1) heat shock transformation, (2) colony selection on selective media, (3) colony picking, and (4) protein expression induction. These steps are implemented as four modular, self-contained OT-2 protocols, allowing users to execute either complete workflows or individual steps as needed (Fig. 1b). Each module is configured through two files: a JavaScript Object Notation (JSON) file specifying OT-2 hardware/labware setup and protocol-specific parameters, and a Comma Separated Value (CSV) file detailing liquid handling instructions such as source wells and transfer volumes. These configurations integrate into a master Python protocol (OT-2 APIs v2.16) that loads directly into the Opentrons app without requiring additional coding.

To facilitate preparation work, APEX provides comprehensive PDF documentation with step-by-step instructions and R-generated visual guides depicting plate contents (Fig. 1c). The entire workflow is packaged as a Nextflow [22] pipeline, with all dependencies contained within a Docker/Singularity image [23], ensuring robust and reproducible software execution. APEX and its detailed documentation are available at: https://github.com/stracquadaniolab/apex-nf and Supplementary File 1, which provides a comprehensive manual with step-by-step instructions for all protocols, including configuration parameters and workflow details.

### Protocol 1 - Heat shock transformation

The most widely used method for delivering exogenous DNA into *E. coli* is heat shock transformation. In brief, the cells and the exogenous DNA are preincubated together in ice-cold calcium chloride. A sudden, rapid shift in temperature enables the uptake of the exogenous DNA. The cells are subsequently recovered in a rich recovery medium such as the Super Optimal broth with Catabolite repression (SOC) medium [24]. The necessary step of this process is ensuring rapid temperature changes. Although the OT-2 temperature module can reach required transformation temperatures, its warm-up and cool-down rates are not sufficiently fast [25], and require additional manual handling, such as adding plate covers to prevent evaporation. Therefore, APEX uses a thermocycler module that enables rapid temperature gradients. By default, cells are incubated at 4 ^°^C for 30 minutes, heat shocked at 42 ^°^C for 30 seconds, and recovered with SOC medium at 37 ^°^C for 1 hour. Users can easily modify these parameters in the configuration files to suit their specific requirements (Supplementary File 1, Protocol 1)

### Protocol 2 - Colony selection

Following transformation, cells are plated on a selective solid medium, typically Lysogeny-Broth (LB) agar supplemented with antibiotic. This ensures that only cells carrying the DNA payload will survive. This traditionally time-consuming step can be efficiently automated using liquid handling robots.

APEX Protocol 2 enables the OT-2, which lacks sensing capabilities, to dispense transformed cells onto agar plates by calculating agar height from the plate base area, weight, and agar density (Fig. 2a). The protocol includes adjustable parameters for customising spotting height relative to the calculated agar height and optional cell dilution before spotting ((Fig. 2b), Supplementary File 1, Protocol 2). APEX supports Thermo Scientific Nunc OmniTray plates (96 transformations in an 8×12 grid) and standard 90mm Petri dishes (32 transformations in an 8×4 grid) with a 3D-printed adapter.

**Figure 2:**
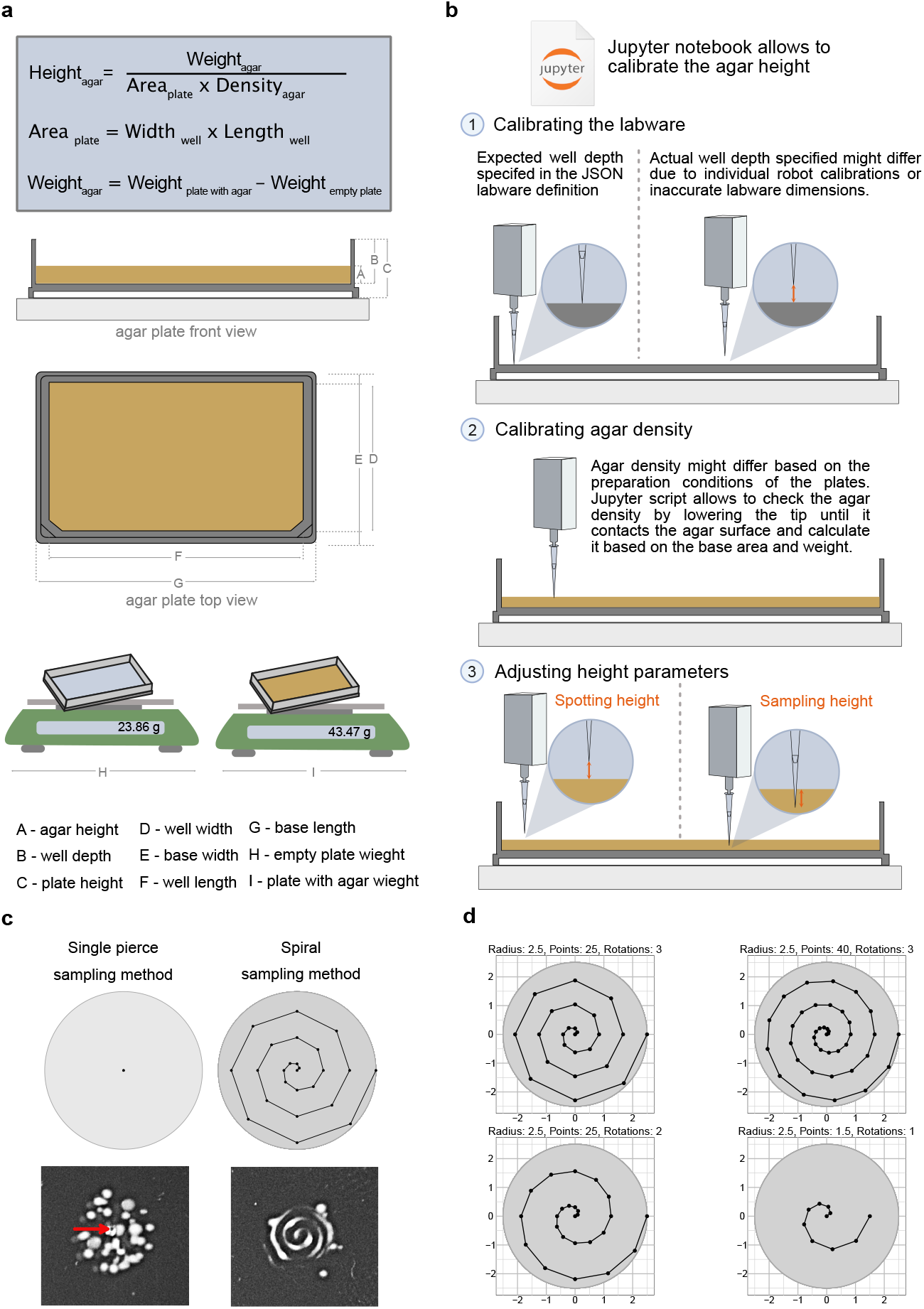
Developing methods for automated agar plate spotting and colony sampling. **a**. Agar height calculation for accurate spotting uses four input parameters: plate base area, agar density, empty plate weight, and plate weight with agar. The height is calculated by determining the agar weight and dividing by the product of base area and agar density. **b**. A Jupyter notebook enables users to verify and adjust agar heights, accounting for variations in robot calibration, plate dimensions, and agar solidification conditions. **c**. Two colony sampling strategies were developed: (i) a single pierce method that targets the center of the initial cell dispersion spot, and (ii) a spiral method where the pipette moves outward in a spiral pattern from the same central coordinates. **d**. The spiral sampling method is customisable through three parameters: radius controls the spiral’s outward reach, number of points determines path resolution, and rotation count defines the number of complete turns.

Agar density was experimentally determined using an interactive, pre-installed Jupyter notebook on the OT-2 to measure agar height via a pipette tip (detailed methodology in Supplementary Materials). Measurements revealed densities of 0.911 g cm^−3^ for rectangular plates and 0.954 g cm^−3^ for round plates (Supplementary Tables 5 and 6). The Jupyter notebook (Supplementary File 2) was further developed to enable users to verify and adjust agar density and spotting height to accommodate variations in plate types, preparation, or individual robot calibrations.

### Protocol 3 - Colony sampling

Selecting single colonies for downstream culturing and protein expression is usually a tedious operation. This step can be automated through the use of specialised equipment, such as the QPix system, which have detection capabilities. However, when the objective is to express or screen proteins in high-throughput, the requirement for single colonies can be relaxed, and non isogenic cultures can be used. In this scenario, we were able to automate this process by using pipette with attached tips to sample colonies from agar plates.

For accurate colony sampling, APEX calculates the agar height using plate measurements (Fig. 2a), with a verification method available to account for any variations in agar height (Fig. 2b). Given that the success rate of this operation depends on the abundance and distribution of colonies across the spot, we developed two strategies (Fig. 2c): (i) a single pierce sampling method for high cell densities, where the tip directly targets the centre of the initial cell dispension, and (ii) a spiral sampling method, where the pipette traces a spiral motion from the same centre coordinates outward for low or uneven cell densities. The spiral method can be customised by adjusting parameters such as radius and number of rotations (Fig. 2d), using slow, smooth movements to minimise disturbance to the agar. Albeit slower, this method improves the likelihood of successful sampling colonies compared to the quicker single pierce method. After sampling, APEX inoculates cells into fresh LB-antibiotic by mixing media in a 96 deep well plate (Supplementary File 1, Protocol 3)

To ensure the propagation of positive transformants and mitigate potential issues with plasmid retention, we implemented an automated colony PCR sample preparation protocol as a quality control measure (Supplementary File 1, Protocol 5). This protocol complements the primary workflow and confirms the presence of the target plasmid in the selected colonies. APEX begins by distributing water into a 96-well PCR plate. Individual colonies are manually picked and resuspended into each well, followed by the automated addition of the PCR master mix. PCR amplification is performed using the Opentrons thermocycler and the PCR products are analysed by agarose gel electrophoresis.

### Protocol 4 - Protein expression induction

After successful transformation and overnight culturing, cells are primed for protein expression induction. This stage enables translation of the target gene into the corresponding protein. The induction method depends on the plasmid vector and expression strain used, typically requiring the addition of chemical inducers such as isopropyl-*β*-D-thiogalactoside (IPTG) or L-arabinose. APEX propagates cells in fresh LB-antibiotic media using an aliquot of the overnight culture. The plate with subculture is then manually transferred to a shaking incubator until cells reach mid-log phase. Following this, the robot dispenses the chemical inducer and the cultures are incubated to produce protein (Supplementary File 1, Protocol 4). To monitor cell growth, we developed a protocol that automates the preparation of samples for Optical Density at 600 nm (OD_600_) measurements in a microplate (Supplementary File 1, Protocol 7).

For protein expression analysis, APEX offers an additional protocol which automates SDS-PAGE sample preparation (Supplementary File 1, Protocol 6). APEX aliquots specified culture volumes into a fresh plate allowing for growth-normalised sampling, which is then processed in an external centrifuge to pellet the cells. The robot removes the supernatant, adds and mixes lysis solution with the pellet. After incubation, the plate is again centrifuged, and APEX transfers the supernatant to a microplate where SDS-PAGE sample buffer is added. The samples are then heat-denatured in the thermocycler, producing cell-free protein extracts ready for analysis.

### Optimisation of transformation volume

When adapting laboratory protocols for automation, volumes are typically miniaturised to enhance efficiency and reduce costs. To evaluate the effects of volume reduction on Transformation Efficiency (TE), standard volumes recommended by the competent cell supplier were adjusted to fit a 200 µl PCR plate suitable for a thermocycler module. We evaluated three transformation conditions: High Volume (HV), Medium Volume (MV), and Low Volume (LV) (Table 1). Using Protocol 1, we performed automated heat shock transformations in DH5*α* cells using plasmids pEX05 (pJKR-H-araC encoding sfGFP, 1.5 *×* 10^−4^ pmol) and pEX01 (pUC19 as positive control, 1.5 *×* 10^−5^ pmol) (Fig. 3a).

**Table 1:** Configurations of miniaturised transformation volumes. Volume of DNA, cells, and SOC medium used for High Volume (HV), Medium Volume (MV), and Low Volume (LV) configurations.

| Configuration | DNA volume (µl) | Cells volume (µl) | SOC volume (µl) | Total volume (µl) |
| --- | --- | --- | --- | --- |
| HV | 3 | 30 | 150 | 183 |
| ML | 2 | 20 | 100 | 122 |
| LV | 1 | 10 | 50 | 61 |

**Figure 3:**
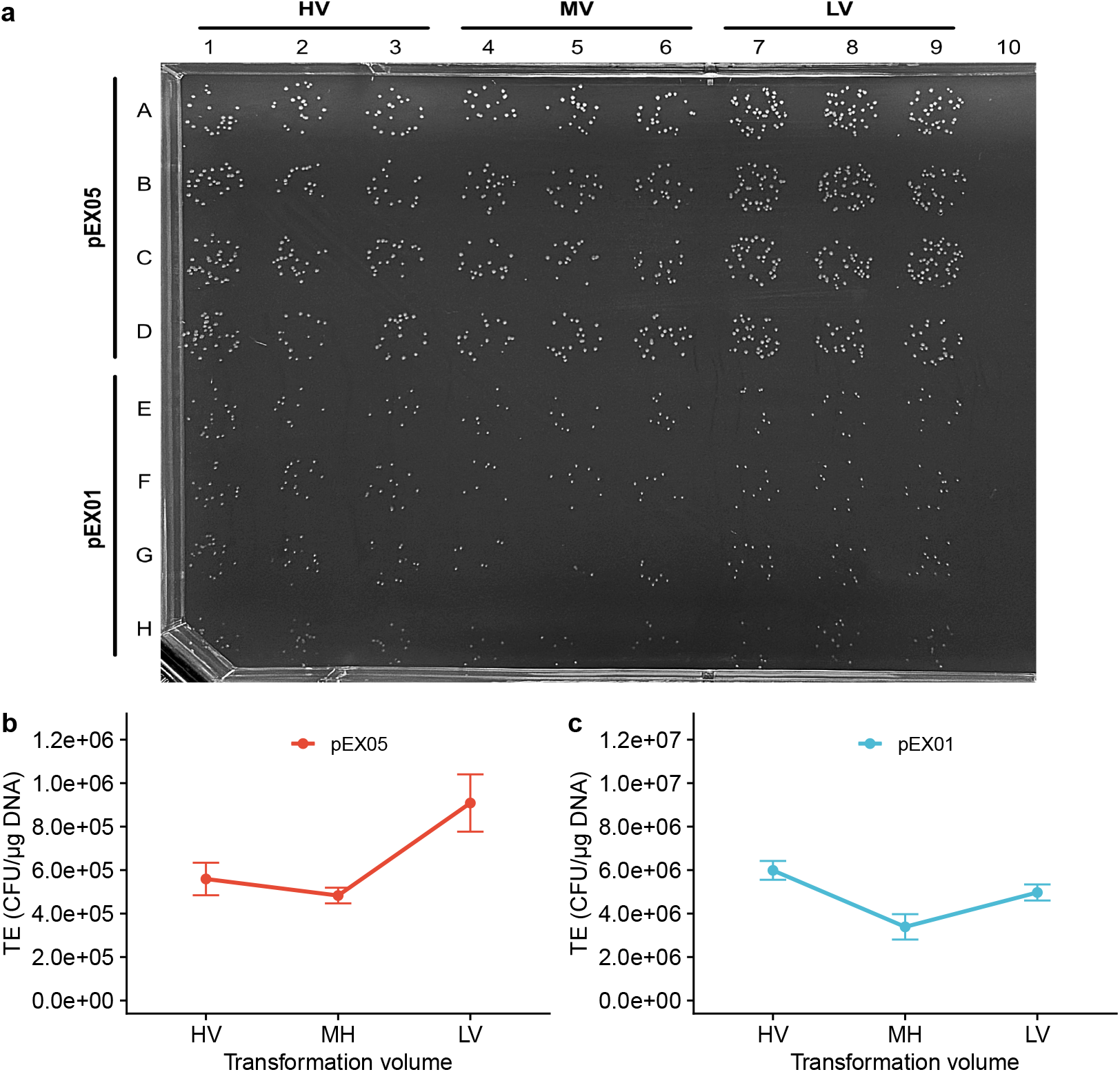
Transformation Efficiency (TE) of *E. coli* DH5*α* at reduced volumes. **a**. DH5*α* transformations with pEX05 (Rows A-D) and pEX01 (Rows E-H). Columns 1-3 show High Volume (HV) transformations, columns 4-6 Medium Volume (MV), and columns 7-9 Low Volume (LV), with each row representing a technical replicate. Row 10 is a negative control. Panels **b** and **c** show TEs (CFU/µg) of pEX05 and pEX01, respectively. Error bars represent standard deviation from the mean value (n=4).

For plasmid pEX05, the LV configuration achieved the highest TE, showing 88% and 63% increases compared to MV and HV configurations, respectively (Fig. 3b, Supplementary Table 2). For plasmid pEX01, the HV configuration yielded the highest TE, showing 77% and 21% increases compared to MV and LV configurations (Fig. 3c, Supplementary Table 2). While TEs varied between plasmids and volumes, all conditions maintained robust TEs. We selected the LV configuration for subsequent experiments, as it reduces reagent consumption while maintaining high TE and enables higher throughput through faster processing of smaller volumes.

### Evaluating APEX and manual TE using different plasmid sizes

It is known that *E. coli* TE generally decreases with increasing plasmid size [24]. Thus, to evaluate the adaptability of APEX protocols 1 and 2, we challenged these workflows with plasmids ranging from kb to 17.7 kb, using equimolar concentrations (1.5 *×* 10^−4^ pmol) of each plasmid. We transformed DH5*α* cells with: pEX01 (2.7 kb), pEX02 (3.9 kb), pEX03 (4.3 kb), pEX04 (4.7 kb), pEX05 (4.9 kb), pEX06 (7.1 kb), pEX07 (11.6 kb), and pEX08 (17.7 kb). Automated transformations were performed under LV conditions specified by Protocol 1, while manual transformations followed the supplier’s protocol. After SOC recovery, cells were handled as per Protocol 2. For spotting, pEX01 and pEX02 required four-fold and two-fold dilutions respectively, due to their high TE (see Methods).

All plasmids yielded successful colony formation (Fig. 4a). We compared TEs between automated (Fig. 4b) and manual (Fig. 4c) methods (Supplementary Table 3). Statistical analysis revealed no significant differences between automated and manual transformation methods (Supplementary Table 4), while both methods showed the expected inverse relationship between plasmid size and TE. These results demonstrate that APEX achieves comparable TEs to manual methods while offering increased throughput.

**Figure 4:**
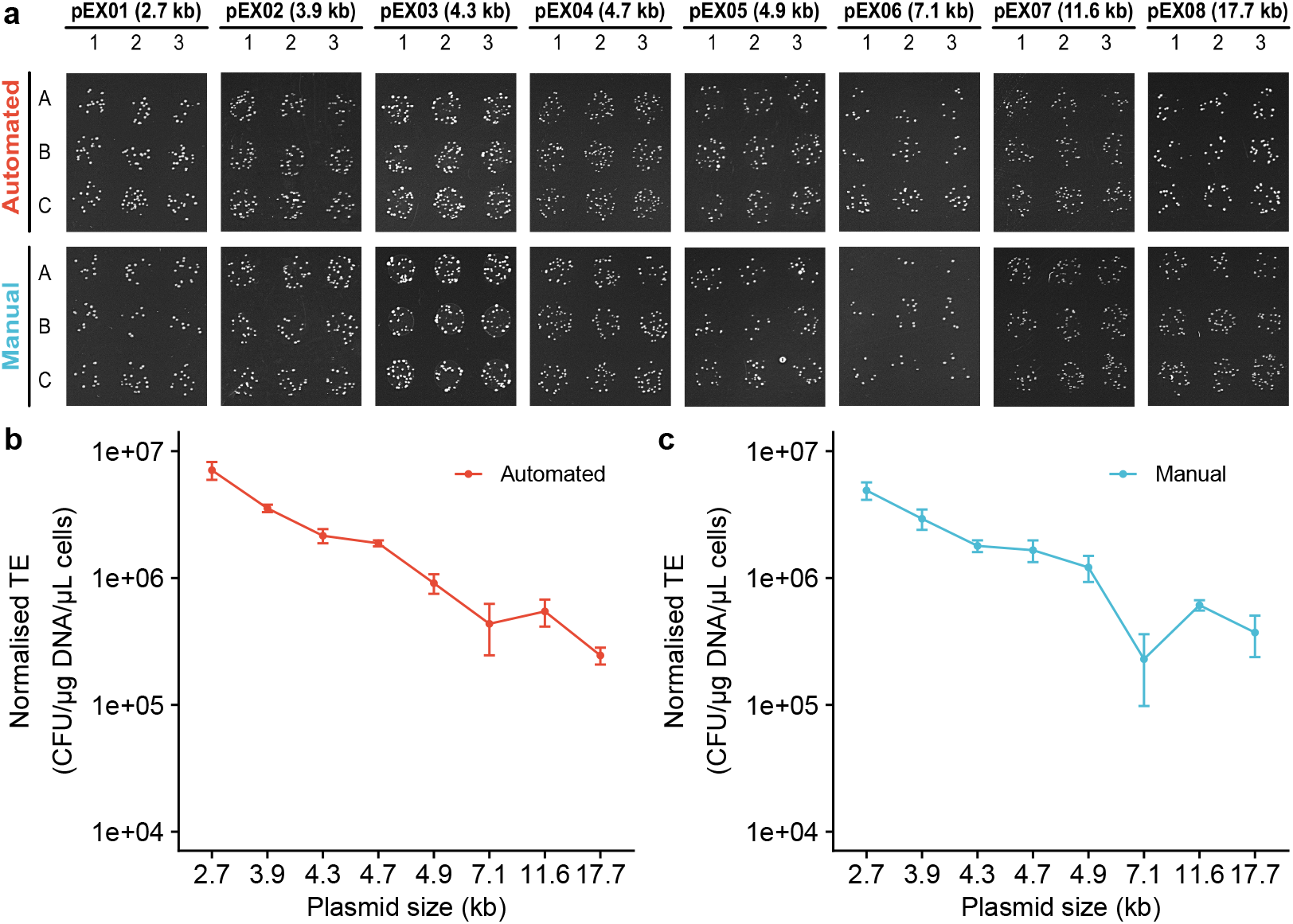
Comparison of automated versus manual transformation methods in DH5*α* across plasmid sizes. **a**. Transformation of DH5*α* cells with eight plasmids ranging from 2.7 kb to 17.7 kb (pEX01-pEX08). All transformations used 1 µl of DNA at 1.5 *×* 10^−4^ pmol. APEXProtocol 1 used reduced volumes (LV configuration, 10 µl cells, 50 µl SOC), while the manual method followed manufacturer’s protocol (50 µl cells, 250 µl SOC). Each condition tested in biological (Rows A-C) and technical (Columns 1-3) triplicates. Spotted cells (5 µl) were plated on antibiotic-supplemented LB agar, with pEX01 and pEX02 diluted 4-fold and 2-fold respectively before spotting. **b** and **c** show the normalised TE for APEX and manual methods, respectively. Error bars show standard deviation (n=3).

### Validation of colony sampling, microculturing and protein expression protocols

To evaluate the end-to-end capability of the APEX workflow, we used sfGFP as our recombinant protein target. We performed 32 transformations of the pEX05 plasmid in BL21(DE3) cells using Protocol 1, followed by colony selection using both undiluted and five-fold diluted outgrowth, as per Protocol 2. This setup created variable colony densities to assess two colony sampling strategies: (i) high density for the single pierce method and (ii) low density for the spiral method. All transformations yielded colonies with the intended cell densities (Fig. 5a). OD_600_ measurements from overnight cultures confirmed successful cell propagation (Fig. 5b).

**Figure 5:**
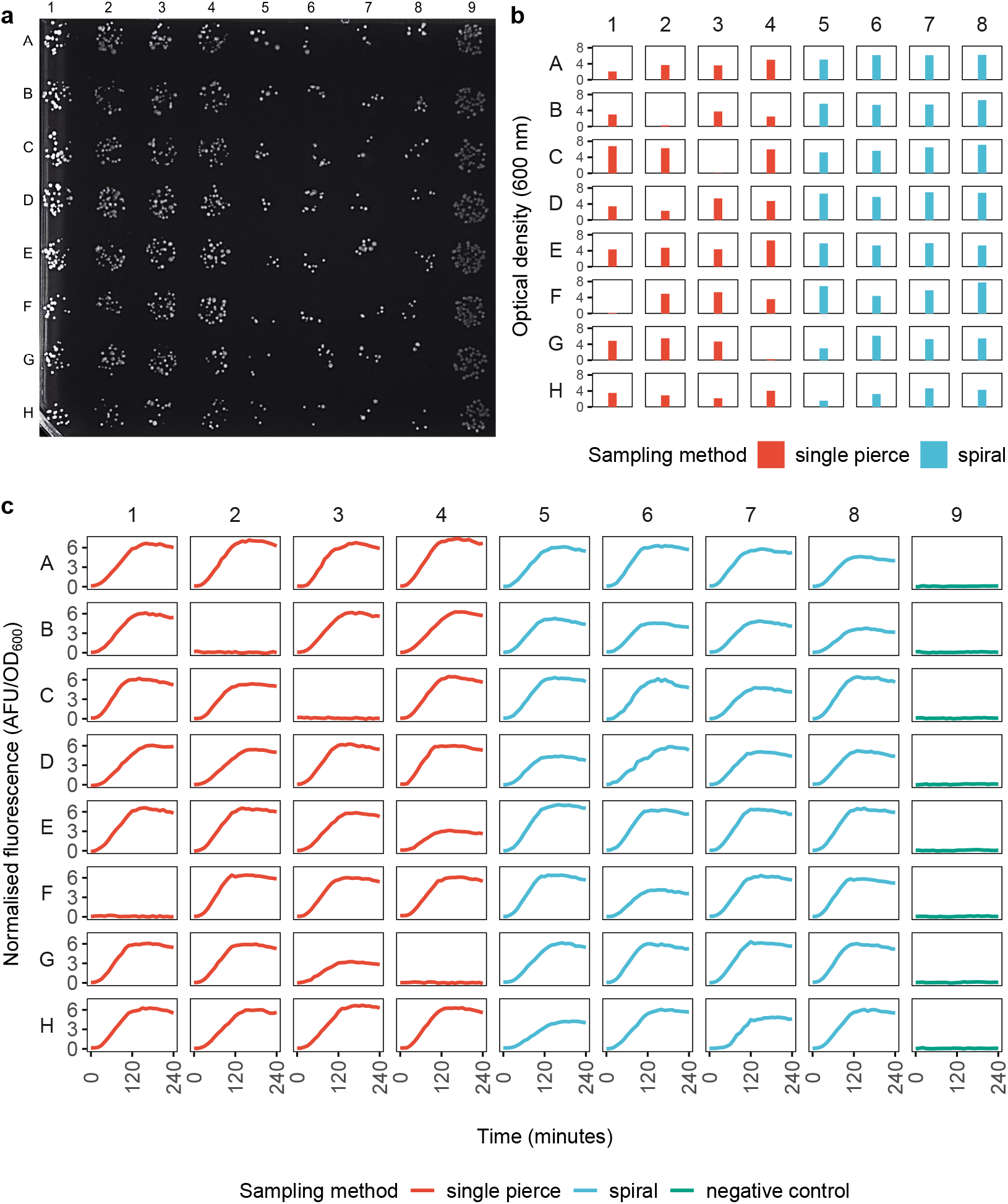
Evaluation of automated colony sampling methods for sfGFP expression in *E. coli* BL21(DE3). **a**. *E. coli* BL21(DE3) cells transformed (Protocol 1) with plasmids pEX05 encoding sfGFP (Columns 1-8), and pEX01 (Column 9, positive control). Cells were spotted directly (Columns 1-4) or diluted five-fold (Columns 5-8) onto selective agar to generate high- and low-density colonies respectively (Protocol 2). **b**. Growth comparison (OD_600_) of overnight cultures sampled using Protocol 3’s two methods: single pierce sampling from high-density colonies (Columns 1-4) and spiral sampling from low-density colonies (Columns 5-8). **c**. sfGFP expression analysis (Protocol 4) showing normalised fluorescence (AFU/OD_600_) over 240 minutes following L-arabinose induction (1.33 mM, Columns 1-8). Column 9 (no inducer) serves as negative control.

The single pierce method successfully sampled and cultivated cells in 28 out of 32 biological replicates, with reduced success rates when colonies were non-centrally distributed. The spiral method achieved successful sampling across all biological replicates, demonstrating its robustness. Subcultures were induced with L-arabinose to express sfGFP (Protocol 4), with fluorescence measurements confirming protein production in all cultures (Fig. 5c).

To screen for the presence of the target plasmid, we used the automated colony PCR sample preparation protocol. Eight transformations were performed, with three colonies from each spot manually selected for automated PCR processing. All colonies tested positive for the sfGFP insert (Supplementary Fig. 1), further supporting the reliability of our pipeline. All together, these results demonstrate the robust performance of the APEX workflow for automated colony processing and protein expression in *E. coli*.

### Analysis of the end-to-end protein production across diverse protein targets

To demonstrate the versatility of the APEX pipeline for heterologous protein production, we selected three enzymes from distinct prokaryotic organisms, covering a range of molecular weights: 4’-phosphopantetheinyl transferase Sfp from *Bacillus subtilis* (29 kDa, pEX04) [27], *ω*-transaminase CVTA from *Chromobacterium violaceum* (49 kDa, pEX09) [28], and an engineered polyketide synthase module from *Mycobacterium tuberculosis* (222 kDa, pEX07) [29]. The complete APEX workflow — including transformation, colony selection/sampling, microculturing and IPTG-inducible protein expression — was performed using BL21(DE3) cells and compared with manual processing. For both methods, protein expression was conducted at 20 ^°^C at two microculture volumes (0.5 mL and 1 mL) in a 96 deep well plate. SDS-PAGE sample preparation was automated for the APEX workflow, while manual workflow samples were processed by hand.

While a slight growth deficit was observed in 1 mL microcultures compared to 0.5 mL microcultures (Fig. 6a, 6b), likely due to reduced oxygen transfer, all three target proteins were successfully expressed and remained soluble in both automated and manual workflows at both culture volumes (Fig. 6c, 6d). These results further demonstrate the versatility and reliability of the APEX platform for automating heterologous protein expression in *E. coli* across a range of protein sizes and microculturing conditions.

**Figure 6:**
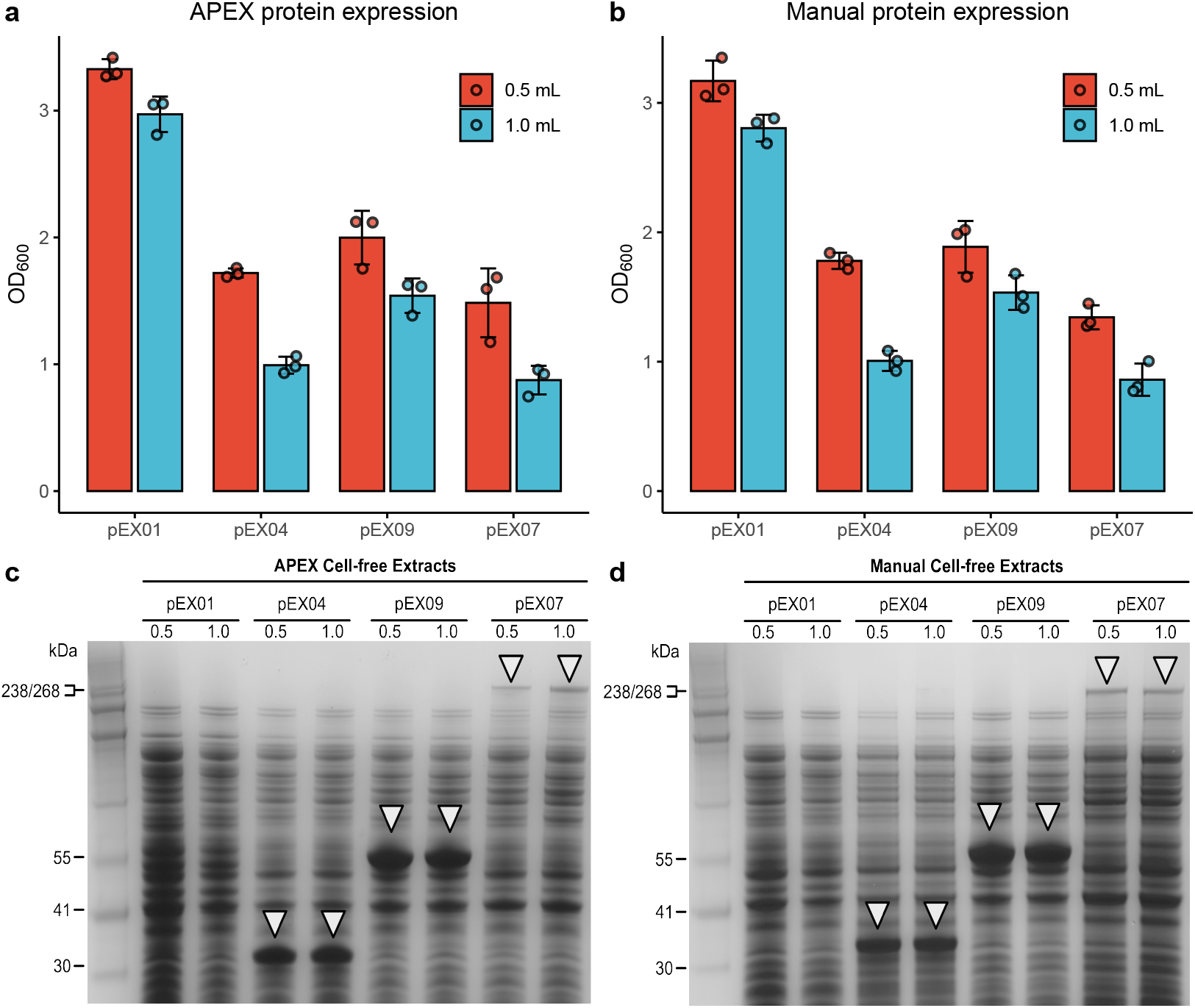
Comparison of APEX and manual protein expression methods at different protein sizes and microculture volumes. **a-b**. Cell density (OD_600_) comparison after 18-hour expression at 20 ^°^C in *E. coli* BL21(DE3). Cells transformed with plasmids encoding proteins of varying sizes (pEX01 control, pEX04 [29 kDa], pEX09 [49 kDa], and pEX07 [222 kDa]) were grown in 96-deep V-well plates at two volumes: 0.5 mL and 1 mL. Results shown for APEX (**a**) and manual (**b**) workflows. **c-d**. SDS-PAGE analysis of soluble protein fractions from the corresponding cultures. Samples from IPTG-induced (0.1 mM) cultures at both 0.5 mL and 1 mL scales were analysed using automated (**c**) and manual (**d**) methods. Target proteins (white triangles) were verified against HiMark pre-stained protein standard.

## Discussion

Conventional microbiological techniques and methodologies are often not amenable to high-throughput automation. To address this bottleneck, we developed an open-source, highly-customisable suite of protocols for automated microbial handling and recombinant protein expression, named APEX. APEX interfaces with the affordable Opentrons OT-2 liquid handling robot, using a minimal hardware setup. APEX can account for different hardware configurations and requirements as specified by the operator, and can be challenged to transform different *E. coli* cell lines with a variety of expression plasmids, encoding a diverse array of protein targets. We demonstrate the time and resource advantages of APEX-programmed automation and microculturing without compromising TE or soluble protein expression.

APEX was designed to deliver seamless end-to-end protocols while minimising user intervention.

Unlike existing solutions that often require programming skills, APEX’s spreadsheet-based configuration enables high customisability without coding knowledge. Current approaches frequently suffer from scattered protocol components that hinder reproducibility or complex implementations requiring programming for customisation. In contrast, APEX consolidates all components within a unified pipeline with a detailed step-by-step documentation, making protocol creation straightforward and reproducible across different laboratory environments. Overall, APEX offers a robust platform for automated *E. coli* cultivation and recombinant protein production; we anticipate that these user-friendly features combined with APEX’s versatility will extend the accessibility of robust, high-throughput automation to small and medium-sized laboratories.

While APEX provides a robust and accessible workflow for protein expression, it has limitations. First, our colony sampling protocol does not guarantee the selection of isogenic clones due to the lack of imaging capabilities of the OT-2, which may be a concern when dealing with issues such as plasmid retention or toxicity. For high-throughout screening/workflows, this may not be a stringent requirement when performed alongside biological or technical replication, at the user’s discretion. Nevertheless, the cumulative time and resources saved using the reamining APEX protocols still offers a substantial benefit over manual handling.

Moreover, the absence of OT-2 compatible centrifuges inherently introduces manual samples processing steps. While integrated centrifugation is available in high-end automation systems typically found in specialised facilities like biofoundries, our focus remains on providing accessible automation solutions for standard laboratory settings.

Taken together, APEX offers a robust platform for automated *E. coli* cultivation and recombinant protein production. By combining open-source accessibility with reliable performance, we anticipate that APEX will enable small and medium-sized laboratories to implement high-throughput automation in their research workflows.

## Materials and methods

### Reagents, Strains and Plasmids

Two *E. coli* strains were used: DH5*α* and BL21(DE3), both sourced from Thermo Fisher Scientific. An overview of plasmids used in this study can be found in Table 2. pUC19 was purchased from New England Biolabs and pJKR-H-araC [30] was a gift from Prof. George Church (Addgene plasmid # 62563). Plasmids pEX04, pEX06, pEX07, pEX08, pEX09 were kindly gifted by Prof. Dominic Campopiano at the University of Edinburgh. Plasmids were prepared using the GeneJET Plasmid Miniprep Kit (Thermo Fisher Scientific) following the supplier’s protocol. Antibiotics were sourced as follows: carbenicillin from Thermo BioReagents, kanamycin from Gibco, zeocin and chloramphenicol from Thermo Fisher Scientific, and chloramphenicol, used at concentrations of 100 µg ml^−1^, 50 µg ml^−1^, 25 µg ml^−1^, and 25 µg ml^−1^ respectively.

**Table 2.** List of plasmids used for APEX experimental validation.

| ID | Size (bp) | Vector | Gene Product | Origin | Resistance |
| --- | --- | --- | --- | --- | --- |
| pEX01 | 2686 | pUC19 | Empty | N/A | AmpR |
| pEX02 | 3943 | pJUMP27-1A(sfGFP) | N-acetyl-L-glutamate Kinase | *Escherichia coli* | KanR |
| pEX03 | 4287 | pGAPZα B | α-Galactosidase | *Homo sapiens* | BleoR |
| pEX04 | 4695 | pACYC-Duet | 4'-phosphopantetheinyl Transferase | *Bacillus subtilis* | CmR |
| pEX05 | 4897 | pJKR-H-araC | Superfolder Green Fluorescent Protein | *Aequorea victoria* | AmpR |
| pEX06 | 7109 | pET26b | Fatty Acyl-ACP Ligase | *Corynebacterium glutamicum* | KanR |
| pEX07 | 11610 | pET26b | Engineered Polyketide Synthase Module | *Mycobacterium tuberculosis* | KanR |
| pEX08 | 17658 | pET26b | Polyketide Synthase Bimodule | *Mycobacterium tuberculosis* | KanR |
| pEX09 | 6568 | pET28a | ω-Transaminase | *Chromobacterium violaceum* | KanR |

Lysogeny Broth (LB) was prepared with 5 g l^−1^ yeast extract, 10 g l^−1^ tryptone, and 10 g l^−1^ NaCl, all obtained from Merck Millipore. Super Optimal Broth with Catabolite Repression (SOC) was prepared with 2.5 g l^−1^ yeast extract, 10 g l^−1^ tryptone, and 10 mmol l^−1^ NaCl, 10 mmol l^−1^ KCl and 20 mmol l^−1^ glucose (both from Thermo Fisher Scientific), and 10 mmol l^−1^ MgSO_4_ and 10 mmol l^−1^ MgCl_2_ (both from Sigma-Aldrich). LB agar was prepared with 5 g l^−1^ yeast extract, 10 g l^−1^ tryptone, 10 g l^−1^ NaCl, and 1.5 % agar (Oxoid). For labware and hardware used in this study see Supplementary Table 1.

### Competent cells preparation

A single colony of *E. coli* strain was inoculated into 5 mL of LB medium and incubated overnight at 37 ^°^C with shaking. The overnight culture was 100-fold diluted into 50 mL of fresh LB medium to grow at 37 ^°^C and 300 rpm until it reached OD_600_ = 0.4. The cells were incubated on ice for 20 minutes and harvested by centrifugation at 4000 *×*g and 4 ^°^C for 10 minutes. Cells were prepared as calcium-competent using the Hanahan method [31], and the pellets were resuspended in 2 mL of 0.1 M CaCl_2_ with 15% (v/v) glycerol solution and stored at −80 ^°^C.

### Calculation of TE

Transformation Efficiency (TE) is quantified as the number of colony-forming units (CFUs) per µg of plasmid DNA. To ensure consistency, the following parameters were standardised: resuspension volume at 2 mL from 50 mL bacterial cultures, cell density at OD_600_ = 0.4, use of the same bacterial stock within each experiment, same heat shock conditions, and plating 5 µL of transformation. Images of the agar plates were taken using the Syngene™ NuGenius Gel Documentation System and CFUs was counted using ImageJ [32]. TE was calculated as follows:

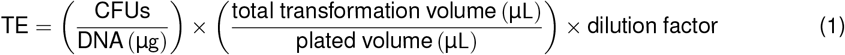

Moreover, to account for variations in aliquots of competent cells and the cells-to-DNA ratio, TE was normalised as:

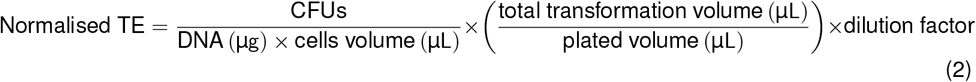

### super folder Green Fluorescent Protein (sfGFP) experiment

Overnight cultures of *E. coli* BL21(DE3) transformed with the pJKR-H-araC plasmid were diluted and grown to an OD_600_ of approximately 0.2, at which point they were induced with 1.33 mM L-arabinose. 150 µL of cells were transferred to a clear-bottom 96-well black plate (Thermo Fisher Scientific) and incubated with shaking at 37 ^°^C and 960 rpm high speed in a Varioskan LUX multimode microplate reader (Thermo Fisher Scientific). OD_600_ and sfGFP fluorescence (excitation 485 nm, emission 528 nm) were measured every 10 minutes for 4 hours, and fluorescence (AFU) normalised to OD_600_ was reported.

### Protein expression and SDS-PAGE analysis

Expression analysis was performed in BL21(DE3) cells using plasmids pEX01, pEX04, pEX07, and pEX09. Following transformation and colony selection, cultures were back-diluted to OD_600_ = 0.1 in fresh LB media in a 96 V-well 2 mL plate (Greiner). The cells were grown to mid-log phase at 37 ^°^C, after which protein expression was induced by the addition of 0.1 mM IPTG. The microcultures were sealed using a Breathe-Easy sealing membrane (Diversified Biotech) and incubated overnight at 20 ^°^C with rigorous shaking. Approximately 2.4 *×* 10^8^ cells were sampled from each microculture and harvested either by microcentrifugation (17 000 ×g, 10 minutes) for manual processing or plate centrifugation (3.884 ×g, 10 minutes) for automated processing. Cell pellets were lysed in BugBuster Mastermix (25 µL, Merck) at room temperature for 15 minutes. The lysate was clarified using the same centrifugation conditions. A sample of the cell-free extract (20 µL) was diluted 2-fold in standard 2X SDS-PAGE sample buffer and boiled for 6 minutes. The samples were resolved on a NuPAGE 4– 12% Bis-Tris precast gel (Invitrogen) using 1X NuPAGE MES SDS running buffer (200 V, 35 minutes). The gel was recovered and stained using InstantBlue Coomassie protein stain (Abcam).

## Contributions

G.S. conceived the study. M.K developed the APEX protocols with help from M.A.H. and supervised by G.S. M.K. and M.A.H. performed lab experiments. All experimental work was supervised by G.S. G.S., M.K. and M.A.H. wrote the manuscript.

## Acknowledgments

This work was supported by the UKRI EPSRC Fellowship (EP/V033794/1) and grant (EP/Y01913X/1), and the IBioIC feasibility funding (FF-2022-04) to G.S, and the UKRI Biotechnology and Biological Sciences Research Council (BBSRC) grant BB/T00875X/1 for M.K. We also thank the Edinburgh Genome Foundry (EGF) for the support and useful discussions, and Maximilian Hingerl for providing external validation of the APEX protocols.

