## Supplementary materials for "APEX: Automated Protein EXpression in *Escherichia coli*"

| Hardware/Labware | Source | Identifier |
|---|---|---|
| Thermocycler Module GEN 1 | Opentrons | N/A |
| 20 µL 8-Channel GEN2 Pipette | Opentrons | 999-00005 |
| 20 µL Single-Channel GEN2 Pipette | Opentrons | 999-00002 |
| 300 µL 8-Channel GEN2 Pipette | Opentrons | 999-00006 |
| 300 µL Single-Channel GEN2 Pipette | Opentrons | 999-00003 |
| 20 µL tips | Opentrons | 999-00007 |
| 300 µL tips | Opentrons | 999-00009 |
| 96-well deep-well plate, V-bottom, 2 mL | Greiner | 780285-FD |
| 96-well PCR Plate, V-bottom, Full Skirt 200 µL | Armadillo | 12640965 |
| Nunc OmniTray Single-Well Plate | Thermo Scientific | 140156 |
| Standard 90mm Petri Dishes | Thermo Scientific | 11309283 |

**Supplementary Table 1:** Labware and hardware used in this study.

| Plasmid | Volume | DNA Conc ($\mu g\,\mu L^{-1}$) | CFU 1 | CFU 2 | CFU 3 | CFU 4 | Mean TE $\pm$ SD (CFU/µg DNA) |
|---|---|---|---|---|---|---|---|
| pEX05 | LV | $2.7 \times 10^{-5}$ | 38 | 39 | 38 | 28 | $908\,646 \pm 131\,865$ |
| | MH | $2.7 \times 10^{-5}$ | 19 | 20 | 17 | 20 | $482\,917 \pm 35\,945$ |
| | HV | $2.7 \times 10^{-5}$ | 21 | 19 | 26 | 22 | $559\,167 \pm 474\,825$ |
| pEX01 | LV | $4.8 \times 10^{-4}$ | 10 | 11 | 11 | 12 | $4\,970\,370 \pm 368\,935$ |
| | MV | $4.8 \times 10^{-4}$ | 9 | 8 | 6 | 7 | $3\,388\,889 \pm 583\,338$ |
| | HV | $4.8 \times 10^{-4}$ | 12 | 14 | 14 | 13 | $5\,987\,037 \pm 432\,615$ |

**Supplementary Table 2:** Transformation Efficiency (TE) of *E. coli* DH5$\alpha$ harboring plasmids pEX05 and pEX01, at low (LV), medium (MV), and high volumes (HV). Results are reported as colony-forming units (CFU) across four biological replicates. Standard deviations (SD) are calculated from the mean TE, based on four replicates (n=4).

| Plasmid | Size (kb) | DNA conc ($\mu g\,\mu L^{-1}$) | CFU 1 | CFU 2 | CFU 3 | Mean TE ± SD (CFU/µg DNA/µL cells) |
|---|---|---|---|---|---|---|
| | | | Automated | | | |
| pEX01 | 2.7 | $2.7 \times 10^{-4}$ | 13 | 16 | 18 | 7079012 ± 1137136 |
| pEX02 | 3.9 | $3.9 \times 10^{-4}$ | 21 | 24 | 23 | 3545299 ± 238921 |
| pEX03 | 4.3 | $4.2 \times 10^{-4}$ | 28 | 27 | 34 | 2154365 ± 274931 |
| pEX04 | 4.7 | $4.6 \times 10^{-4}$ | 28 | 27 | 30 | 1878623 ± 101282 |
| pEX05 | 4.9 | $4.8 \times 10^{-4}$ | 14 | 12 | 17 | 910764 ± 159910 |
| pEX06 | 7.1 | $7.0 \times 10^{-4}$ | 8 | 7 | 15 | 435714 ± 189923 |
| pEX07 | 11.6 | $1.1 \times 10^{-3}$ | 16 | 18 | 25 | 545303 ± 131034 |
| pEX08 | 17.7 | $1.7 \times 10^{-3}$ | 12 | 13 | 16 | 245196 ± 37348 |
| | | | Manual | | | |
| pEX01 | 2.7 | $2.7 \times 10^{-4}$ | 12 | 9 | 12 | 4905185 ± 772366 |
| pEX02 | 3.9 | $3.9 \times 10^{-4}$ | 23 | 17 | 17 | 2932821 ± 534715 |
| pEX03 | 4.3 | $4.2 \times 10^{-4}$ | 26 | 22 | 27 | 1791667 ± 189612 |
| pEX04 | 4.7 | $4.6 \times 10^{-4}$ | 22 | 31 | 23 | 1657681 ± 322782 |
| pEX05 | 4.9 | $4.8 \times 10^{-4}$ | 15 | 24 | 19 | 1212361 ± 282768 |
| pEX06 | 7.1 | $7.0 \times 10^{-4}$ | 2 | 8 | 6 | 229333 ± 131367 |
| pEX07 | 11.6 | $1.1 \times 10^{-3}$ | 20 | 23 | 24 | 611121 ± 56962 |
| pEX08 | 17.7 | $1.7 \times 10^{-3}$ | 13 | 22 | 28 | 371824 ± 133676 |

**Supplementary Table 3:** TE of *E. coli* BL21(DE3) harboring plasmids pEX01, pEX02, pEX03, pEX04, pEX05, pEX06, pEX07. Results are reported as colony-forming units (CFU) across three biological replicates. Standard deviations (SD) are calculated from the mean TE, based on four replicates (n=4).

| Plasmid ID | Plasmid size (kb) | Statistic | P-value | Conf. Interval Low | Conf. Interval High |
|---|---|---|---|---|---|
| pEX01 | 2686 | 2.74 | 0.0598 | -152,939 | 4,500,593 |
| pEX02 | 3943 | 1.81 | 0.175 | -516,488 | 1,741,445 |
| pEX03 | 4287 | 1.88 | 0.142 | -200,401 | 925,798 |
| pEX04 | 4695 | 1.13 | 0.359 | -500,776 | 942,660 |
| pEX05 | 4897 | -1.61 | 0.202 | -881,685 | 278,490 |
| pEX06 | 7109 | 1.55 | 0.205 | -182,691 | 595,453 |
| pEX07 | 11610 | -0.798 | 0.488 | -343,671 | 212,035 |
| pEX08 | 17658 | -1.58 | 0.238 | -430,623 | 177,368 |

**Supplementary Table 4:** Statistical analysis comparing TEs for between automated and manual methods by varying plasmid sizes. Differences in TE were determined by an unpaired t-test with default settings for a two-tailed distribution. Differences were considered statistically significant at p $< 0.05$. The data were represented as means ± standard deviation (n=3). Statistical analyses were performed using R software (version 4.3.1). Welch two-sided t-test comparing TE between automated and manual methods by varying plasmid sizes

### Determining the agar height

The Lysogeny Broth (LB) agar medium was prepared by dissolving $5\,\mathrm{g\,L^{-1}}$ yeast extract, $10\,\mathrm{g\,L^{-1}}$ tryptone, $10\,\mathrm{g\,L^{-1}}$ NaCl, and $1.5\,\%$ bacto-agar in deionized water. The molten media was allowed to cool down to $50\,^{\circ}\mathrm{C}$ in a water bath. This temperature prevented the premature solidification of the agar and ensured even distribution across the plate. Using a serological pipette, the media was distributed into the plates on a leveled bench at the following volumes: for rectangular Nunc OmniTray Single-Well plates - $20\,\mathrm{mL}$, $25\,\mathrm{mL}$, and $30\,\mathrm{mL}$; for round $90\,\mathrm{mm}$ petri dishes - $15\,\mathrm{mL}$, $20\,\mathrm{mL}$, and $25\,\mathrm{mL}$. The molten agar was allowed to solidify at room temperature for $15\,\mathrm{min}$, and the plates were then dried in a flow hood for $40\,\mathrm{min}$. Drying the plates is crucial to prevent the diffusion of the transformation spotting.

Using the analytical balance, before pouring the agar, each empty plate was first measured for its mass ($M_{\text{plate}}$), and then the plates with poured agar were measured ($M_{\text{plate+agar}}$). The mass of the agar was calculated using the following equation:

$$M_{\text{agar}}[\,\mathrm{g}] = M_{\text{plate+agar}}[\,\mathrm{g}] - M_{\text{plate}}[\,\mathrm{g}] \tag{1}$$

The agar height was determined by lowering a P20 single-channel pipette via a Jupyter notebook, with a tip attached, to the surface of the agar as shown in Table 5 for rectangular plates and in Table 6 for round plates.

Heights at each point were recorded, and their average was calculated. Subsequently, average weights and heights for each volume were calculated. The density of the agar ($\rho_{\text{agar}}$) at room temperature was calculated using the formula:

$$\rho_{\text{agar}}\left[\mathrm{g\,cm^{-3}}\right] = \frac{M_{\text{agar}}\,[\mathrm{g}]}{\text{Base area}\left[\mathrm{mm^{2}}\right] \times \rho_{\text{agar}}\left[\mathrm{g\,cm^{-3}}\right]} \tag{2}$$

| Parameter | Plate 1 | Plate 2 | Plate 3 |
|---|---|---|---|
| Volume (mL) | 20 | 25 | 30 |
| Empty Plate Weight (g) | 38.9518 | 39.2415 | 38.9445 |
| Plate with Agar Weight (g) | 57.3196 | 61.3349 | 66.1736 |
| Agar Weight (g) | 18.3678 | 22.0934 | 27.2291 |
| Plate base area ($mm^2$) | $9.4692 \times 10^3$ | | |
| Height (mm) A1 | 2.1 | 2.6 | 3.1 |
| Height (mm) A6 | 2.2 | 2.7 | 3.1 |
| Height (mm) A12 | 2.2 | 2.6 | 3.0 |
| Height (mm) C3 | 2.1 | 2.6 | 3.1 |
| Height (mm) C5 | 2.2 | 2.6 | 3.1 |
| Height (mm) C8 | 2.1 | 2.5 | 3.0 |
| Height (mm) C10 | 2.1 | 2.5 | 3.0 |
| Height (mm) F3 | 2.1 | 2.6 | 3.2 |
| Height (mm) F5 | 2.1 | 2.6 | 3.2 |
| Height (mm) F8 | 2.1 | 2.6 | 3.1 |
| Height (mm) F10 | 2.1 | 2.5 | 3.0 |
| Height (mm) H1 | 2.2 | 2.7 | 3.2 |
| Height (mm) H7 | 2.2 | 2.6 | 3.2 |
| Height (mm) H12 | 2.2 | 2.6 | 3.1 |
| Mean Height (mm) | 2.1 | 2.6 | 3.1 |
| Density ($g\,cm^{-3}$) | 0.905 | 0.900 | 0.928 |
| Average Density ($g\,cm^{-3}$) $\pm$ SD | 0.911 $\pm$ 0.0122 | | |

**Supplementary Table 5:** Agar height measurements used to determine agar density for rectangular plates (Thermo Scientific Nunc OmniTray) across three different weights.

| Parameter | Plate 1 | Plate 2 | Plate 3 |
|---|---|---|---|
| Volume (mL) | 15 | 20 | 25 |
| Empty Plate Weight (g) | 14.2407 | 14.2197 | 14.2559 |
| Plate with Agar Weight (g) | 28.7150 | 33.5899 | 38.3507 |
| Agar Weight (g) | 14.4743 | 19.3702 | 24.0948 |
| Plate base area ($mm^2$) | $1.8749 \times 10^3$ | | |
| Height (mm) A1 | 2.6 | 3.4 | 4.4 |
| Height (mm) A4 | 2.5 | 3.3 | 4.3 |
| Height (mm) C2 | 2.6 | 3.4 | 4.3 |
| Height (mm) C3 | 2.5 | 3.5 | 4.3 |
| Height (mm) F2 | 2.6 | 3.4 | 4.3 |
| Height (mm) F3 | 2.6 | 3.4 | 4.3 |
| Height (mm) H1 | 2.7 | 3.5 | 4.4 |
| Height (mm) H4 | 2.7 | 3.5 | 4.4 |
| Mean Height (mm) | 2.6 | 3.4 | 4.3 |
| Density ($g\,cm^{-3}$) | 0.950 | 0.947 | 0.947 |
| Average Density ($g\,cm^{-3}$) $\pm$ SD | 0.948 $\pm$ 0.0014 | | |

**Supplementary Table 6:** Agar height measurements used to determine agar density for round plates (Thermo Scientific 90 mm petri dish) across three different weights.

### Colony PCR validation

Colony PCR was performed to verify the presence of the insert in transformed DH5$\alpha$ colonies with pEX05 (pJKR-H-araC). PCR setup was automated using the APEX protocol on an OT-2 liquid handling robot (see Supplementary File 1 - Manual Protocol 5: Colony PCR - for details). APEX distributed 11.5 µl water to a 96 well PCR plate, paused for manual colony picking and resuspension, then added 13.5 µl OneTaq Quick-Load 2X Master Mix (NEB M0486) with 0.5 µM primers (Supplementary Table 7). Amplification was carried out in the Opentrons thermocycler using the following conditions: initial denaturation at 94 °C for 2 min; 30 cycles of denaturation at 94 °C for 30 s, annealing at 42 °C for 30 s, and extension at 68 °C for 45 s; followed by a final extension at 68 °C for 5 min. PCR products (5 µl) were analysed by electrophoresis on a 1% agarose gel stained with SYBR Safe in 1 x TAE buffer at 80 V for 60 min. HyperLadder™ 1kb (Meridian Bioscience) was used for size estimation. Gels were visualized using a Syngene NuGenius Gel Documentation System, showing a distinct band at ˜700 bp in lanes 1–24 (Figure 1), corresponding to the expected PCR product size.

| **Primer Name** | **Sequence (5' 3')** | **Target** | **Tm (°C)** | **Amplicon Size (bp)** |
|---|---|---|---|---|
| sfGFP_F | CTGTTCACCGGTGTTGTTCC | sfGFP insert | 59.34 | 711 |
| sfGFP_R | GTACAGTTCGTCCATACCGTG | sfGFP insert | 58.48 | 711 |

**Supplementary Table 7:** Primers used for colony PCR validation of sfGFP insert in APEX protocol.

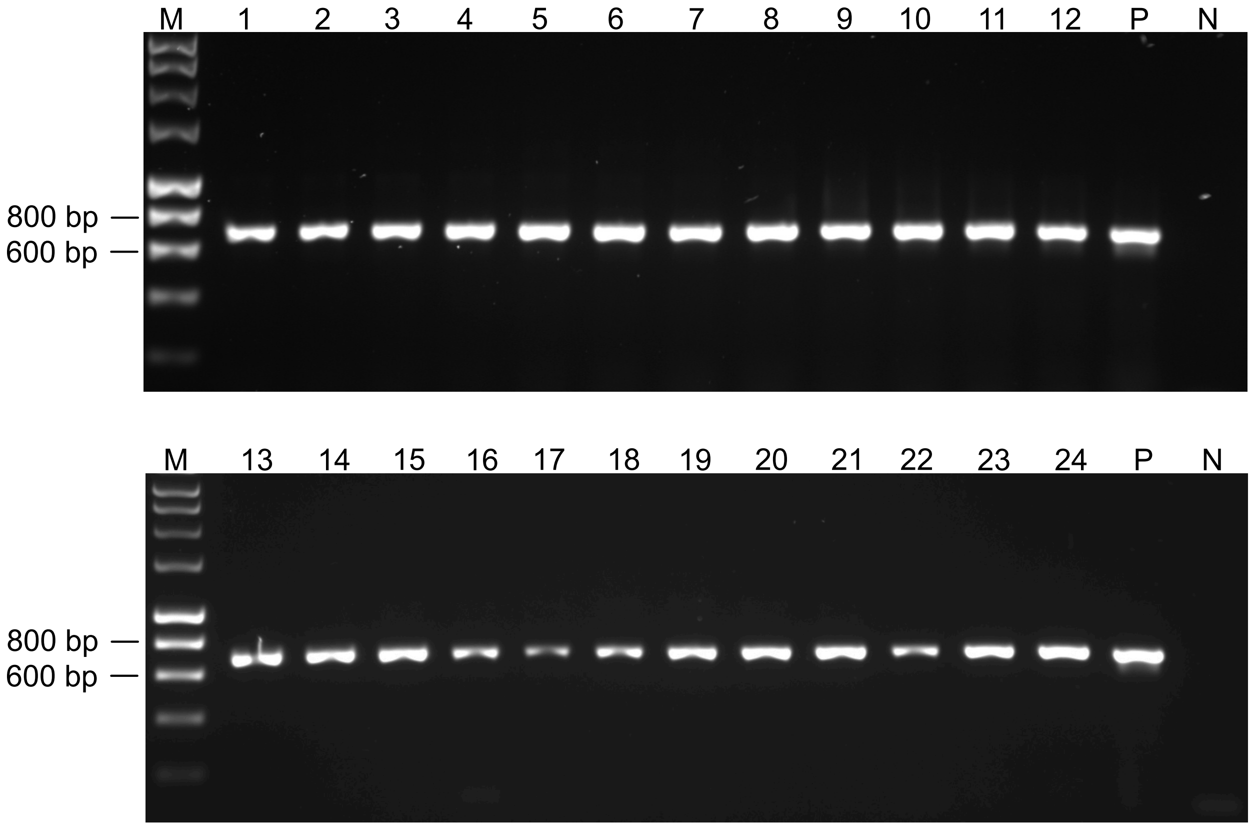


**Figure 1: Automated sample preparation for colony PCR screening demonstartes consistent amplification.** Colony PCR was performed using an automated sample preparation protocol on BL21(DE3) *E. coli* colonies transformed with pJKR-H-araC. Primers were designed to amplify the full-length sfGFP insert (711 bp). PCR products were analyzed on a 1% agarose gel and visualised using SBRsafe DNA stain. Individual colony PCR products are shown in lanes 1-24, with a negative control (no plasmid, lane N) and positive control (lane P). Size reference bands from Meridian Bioscience HyperLadder 1kb are shown on the left. This method enables rapid, parallel screening of multiple colonies with minimal manual intervention.
